# Inhibition of auxin biosynthesis in early rice grains leads to extensive post-fertilisation grain abortion

**DOI:** 10.1101/2023.06.29.547002

**Authors:** Mafroz A. Basunia, Heather M. Nonhebel, David Backhouse, Mary McMillan

## Abstract

In spite of its abundant presence in rice grains, auxin functions during grain development are not understood well. Absence of grain-specific auxin biosynthesis mutants in rice further limits our understanding in this respect. Here, we report a chemical biology approach to inhibit auxin biosynthesis specifically in early rice grains as well as its subsequent effects on final grain yield. Exogenous application of two auxin biosynthesis inhibitors, namely L-kynurenine (L-Kyn) and 4-phenoxyphenylboronic acid (PPBo), to spikelets daily from three to ten days after pollination (DAP) significantly reduced grain levels of indole-3-acetic acid (IAA), the predominant *in planta* auxin. The inhibitor-treated panicles showed extensive post-fertilisation seed abortion, leading to drastic reduction in total panicle weight at maturity. Locally synthesised auxin during early grain development may therefore play a crucial role in grain retention. This chemical biology approach can be an easy and cost-effective way to study auxin biosynthesis and signalling during grain development in rice and possibly other cereal crops.

**Highlight:** Auxin synthesised in early rice grains may play a crucial role in grain retention.

## Introduction

Rice (*Oryza sativa* L.) grains accumulate the largest concentration of free auxin (indole-3-acetic acid, IAA) found anywhere in the plant (Matsuda et al. 2005). Two *Tryptophan Aminotransferase of Arabidopsis1*/*Tryptophan Aminotransferase-Related* (*TAA1/TAR*) genes (*OsTAR1* and *OsTAR2*) and three grain-specific *YUCCA* genes with differing spatio-temporal expression patterns (*OsYUC9, OsYUC11* and *OsYUC12*) contribute to grain IAA pool via indole-3-pyruvic acid (IPyA) pathway (Abu-Zaitoon et al. 2012; Basunia et al. 2021). Of these, *OsYUC12* has a short-lived expression in the aleurone, sub-aleurone and embryo from 3-8 days after pollination (DAP) (Basunia et al. 2021). *OsYUC9* and *OsYUC11* expression occurs at 5-20 DAP, determining to a large extent the grain IAA content (Abu-Zaitoon et al. 2012; Xu et al. 2021). Although there exists correlative and genetic evidence for a role of IAA in starch biosynthesis during grain-fill (Abu-Zaitoon et al. 2012; Xu et al. 2021), IAA function during early grain development is still poorly understood (Basunia and Nonhebel 2019). Final grain size and weight are determined largely by cellular and molecular events occurring during early stages of grain development (Mizutani et al. 2010; Fahy et al. 2018). Investigation into IAA functions during this period is therefore of crucial importance.

Most of the findings regarding IAA involvement in cereal grain development were made possible by grain-specific auxin biosynthesis mutants in maize such as *de18, dek18* and *mn1* (e.g. Bernardi et al. 2012, 2019). Absence of such mutants in rice is a major limiting factor in our understanding of auxin functions during grain development. Furthermore, the presence of three grain-specific rice *YUCCAs* makes knock-out approach difficult, as loss of one *YUCCA* could be compensated by the remaining ones, e.g. reported by Feng et al. 2019 in wild strawberry and Xu et al. 2021 in rice; conversely, knocking out all the *OsYUCCAs* can be lethal (Guo et al. 2020a). One approach to overcome this problem is to inhibit IAA biosynthesis in rice grains during different stages of development by exogenous application of IAA biosynthesis inhibitors. Several potent inhibitors targeting specifically the TAA1/TARs and YUCCAs in the IPyA pathway have been reported (e.g. Soeno et al. 2010; He et al. 2011; Kakei et al. 2015). However, their action has been demonstrated mostly on the vegetative organs and tissues in *Arabidopsis* (e.g. He et al. 2011) and rice (Watanabe et al. 2021). The utility of chemical biology as a potent tool to analyse molecular mechanisms of auxin signalling during grain development in rice and other cereals remains mostly unexplored. In this study, we therefore aimed to investigate the efficacy of IAA biosynthesis inhibitors in inhibiting IAA biosynthesis in early rice grains as well as the effects of this inhibition on final grain yield. Taking into account the expression of *OsYUC12* at 3-8 DAP and the largest increase in grain IAA content between 4-10 DAP (Abu-Zaitoon et al. 2012; Basunia et al. 2021), we applied two potent IAA biosynthesis inhibitors, viz. L-kynurenine (L-Kyn) and 4-phenoxyphenyl boronic acid (PPBo), targeting TAA1/TARs and YUCCAs, respectively, to rice spikelets daily from 3-10 DAP. We measured grain IAA levels at 5 and 10 DAP to monitor the inhibition efficacy. Yield-related data were collected at maturity in order to examine the effects of early IAA deficiency on final grain yield.

## Material and Methods

### Plant material and growing conditions

Rice plants (*O. sativa* L. ssp. *japonica* cv. Reiziq) were grown in cylindrical plastic pots (50 cm × 15 cm; one plant per pot) in a greenhouse under natural light conditions with 30^0^C/18^0^C day/night temperatures. Plants were watered daily and fertilised fortnightly with Aquasol^®^ (2.0 gm/L) until panicle initiation. Panicles in which approximately half of the spikelets reached anthesis were tagged in the afternoon and the date was recorded as the day of pollination; the following day was designated as one (1) day after pollination (DAP) and so on. Tagged panicles were treated with IAA and IAA biosynthesis inhibitors daily from 3 to 10 DAP (see next section). Panicles were harvested at three time points: 5, 10 and 35 DAP. At 5 and 10 DAP, only superior caryopses were collected, weighed, frozen immediately in liquid nitrogen and stored at -80^0^C until further use. The rest of the panicles were harvested at maturity. They were dried in an oven at 50^0^C for a week and yield-related data were then recorded from each.

### Treatment with exogenous IAA and IAA biosynthesis inhibitors

IAA was applied as exogenous auxin, L-Kyn and PPBo as IAA biosynthesis inhibitors (Sigma-Aldrich). Each chemical was dissolved in dimethyl sulfoxide (DMSO) and applied in two concentrations: 50 µM (T1) and 200 µM (T2) for IAA, 100 µM (T3) and 500 µM (T4) for L-Kyn, and 30 µM (T5) and 100 µM (T6) for PPBo. Final DMSO concentration in all treatment solutions was 0.1%. Each treatment also contained 0.01% Tween-20 (Sigma-Aldrich) to allow better penetration of the chemicals into plant tissues (Fernández and Eichert 2009). A treatment consisting of 0.1% DMSO and 0.01% Tween-20 was applied as control. Six plants were used for each treatment. The total of 42 plants were arranged in a completely randomized block design. Six panicles with homogeneity in appearance were chosen from each plant for chemical application after anthesis. Approximately 3.0 mL of a treatment solution was applied directly and gently to all spikelets on a panicle with a calligraphy brush daily in the afternoon from 3 to 10 DAP.

### Extraction and analysis of IAA from developing rice grains

IAA was extracted from immature rice grains at 5 and 10 DAP. Grain samples (80-100 mg) were finely ground in liquid nitrogen. 200 µl of 65% isopropanol/35% 0.2 M imidazole extraction buffer pH 7.0 (Chen et al. 1988) and 5.0 µl of [^13^C_6_]-IAA internal standard (Cambridge Isotope Laboratories Inc.) were added. Total amount of [^13^C_6_]-IAA standard delivered to 5- and 10-DAP samples was 4.5 ng and 83 ng, respectively; these amounts were similar to those of endogenous grain IAA at 5 and 10 DAP (Abu-Zaitoon et al. 2012). Blank samples containing only the [^13^C_6_]-IAA standard went through the entire extraction and quantification process. Samples were extracted on ice for 1 h with occasional shaking. The supernatant was diluted with 2.0 mL deionized water. Sample clean-up was carried out using the solid-phase extraction (SPE) protocol of Barkawi et al. 2008 with some modification. The diluted supernatants were loaded onto Supelco 1 mL, 50 mg amino SPE cartridges (Sigma-Aldrich), which were pre-washed sequentially with 500 µl hexane, 500 µl acetonitrile, 500 µl water, 500 µl 0.2 M pH 7.0 imidazole and 4.5 mL water. The SPE columns were then washed sequentially with 500 µl each of water, hexane, ethyl acetate, acetonitrile, methanol and 400 µl of 0.25% phosphoric acid. IAA was eluted in 1.8 mL of 0.25% phosphoric acid; eluate pH was adjusted to pH 3.0-3.5 by adding 150 µl of 0.1 M pH 6.0 succinate buffer. The eluates were loaded onto Strata-X SPE columns (60 mg/3 mL; Phenomenex), pre-washed with 1 mL hexane, 1 mL methanol and 2 mL water. The columns were washed with 3×1 mL water and IAA was eluted in 1 mL acetonitrile. The final eluates were stored at -20°C until further use. Before quantification, the eluates were dried under nitrogen gas and re-dissolved in 20 µl acetonitrile and 80 µl 0.1 M acetic acid. An LC-MS/MS multiple reaction monitoring (MRM) with heavy isotope labelled internal standards (Liquid Chromatograph Mass Spectrometer-8050 from Shimadzu, Japan with XBridge™ C18 3.5 µm, 2.1×50 mm column from Phenomenex) was used to measure the IAA content. 20% acetonitrile: 80% 0.01 M acetic acid was used as the HPLC solvent at a flow rate of 0.2 mL·min^-1^. The nebulizing, heating and drying gas flow were 3 L·min^-1^, 10 L·min^-1^ and 10 L·min^-1^, respectively. Interface temperature was 300^0^C, desolvation line (DL) was 250^0^C and the heat block temperature was 400^0^C. The interface used a capillary voltage of 4 kV. The mass spectrometer was operated in MRM mode (collision energy, 14.0 eV), transitions from *m/z* 174.10 to 130.10 for [^12^C_6_] and *m/z* 180.20 to 136.15 for [^13^C_6_]. A series of standard mixtures of [^13^C_6_]-IAA and unlabelled IAA in different ratios ranging from 10:1 to 1:10 was also assayed to confirm the accuracy of quantitative analysis. IAA content was calculated as average from four biological replicates of a treatment.

### Statistical analysis

Results are reported as mean ± standard error of mean (SEM). Statistical analyses were carried out by Minitab^®^ statistical software package (https://www.minitab.com). Data were subjected to one-way analysis of variance (ANOVA). In case of significant difference, mean values were compared using Tukey’s honestly significant difference (HSD) test. A level of *P* < 0.05 was considered to be statistically significant.

## Results

### L-Kyn and PPBo application reduced IAA levels in early rice grains

After application of L-Kyn and PPBo to spikelets daily from 3 to 10 DAP, we measured endogenous IAA levels in treated and untreated grains at 5 and 10 DAP. At 5 DAP, L-Kyn and PPBo treatments reduced significantly grain IAA levels compared to the control (Fig. 1A). As a clear distinction between filled and unfilled/aborted superior grains following chemical treatments became manifest at 10 DAP (see next section), we only used the filled superior grains for IAA quantification of grains at 10 DAP. At this time point, all treatments of L-Kyn and PPBo caused a significant reduction in IAA levels (Fig. 1B); the highest reduction was recorded with 500 µM L-kynurenine when compared to the control. However, at 10 DAP, the lower concentration of PPBo produced a larger reduction in IAA levels than the higher concentration. We also treated the panicles with two different concentrations of IAA for comparison with inhibitor treatments (Fig. 1A-B). At 5 DAP, treatment with 200 µM IAA led to a significant increase in grain IAA content. However, at 10 DAP, we recorded a reduction in grain IAA levels with both treatments, with 200 µM IAA treatment causing the largest reduction. Interestingly, at 10 DAP, the effect of 200 µM IAA in reducing grain IAA content was similar to that of 500 µM L-kynurenine and 30 µM PPBo.

**Fig. 1.**
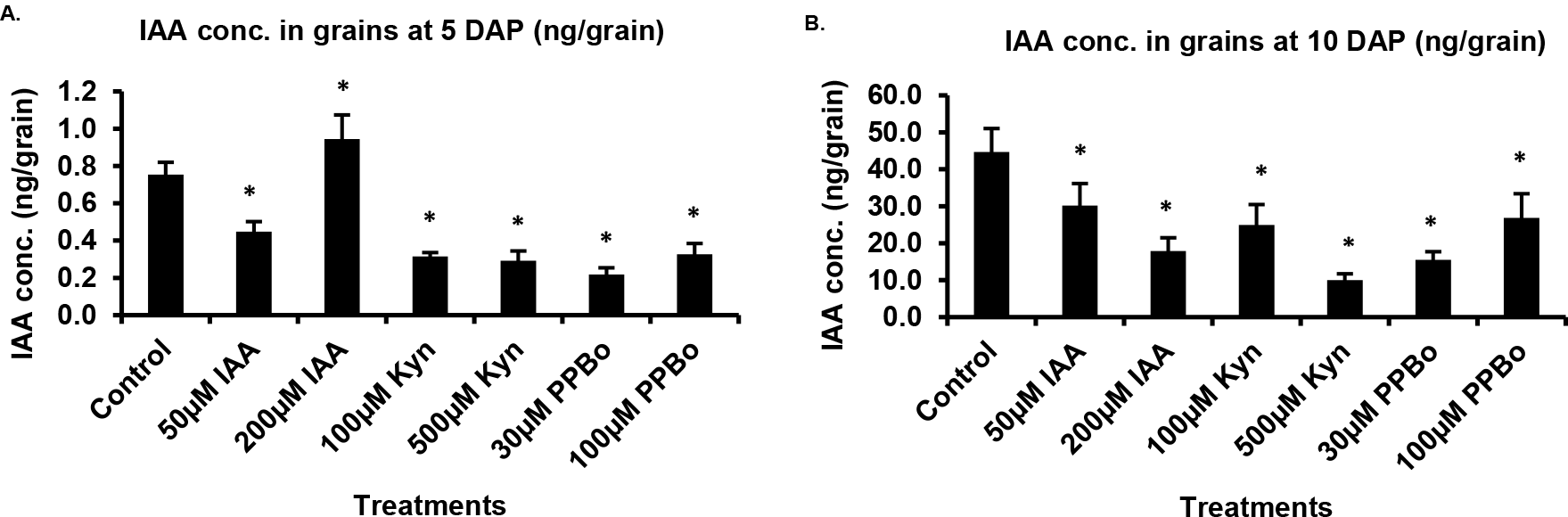
**(A-B)**. Endogenous IAA levels in treated and untreated rice grains at 5 and 10 DAP. IAA and IAA biosynthesis inhibitors (L-kynurenine and PPBo) were applied to spikelets in two different concentrations daily from 3 to 10 DAP. Endogenous IAA content was measured by LC-tandem MS using [^13^C_6_]-IAA as the internal standard. Each data point represents the mean of four biological replicates ± the standard error of the mean (SEM). Data were subjected to one-way analysis of variance (ANOVA). In case of significant difference, mean values were compared using Tukey’s honestly significant difference (HSD) test. Asterisk denotes significant difference of a treatment when compared to the control at *P* <0.05. Kyn= L-kynurenine; PPBo= 4-phenoxyphenylboronic acid; DAP= days after pollination.

### Exogenous application of IAA biosynthesis inhibitors led to seed abortion

Following exogenous application of IAA, L-Kyn and PPBo, we harvested the panicles and recorded yield-related data at three time points: 5, 10 and 35 DAP (at maturity). At 5 DAP, all treatments caused significant reductions in fresh weight of superior grains compared to control (Fig. 2A). At 10 DAP, both concentrations of L-Kyn caused a reduction in grain weight. However, only 200 µM IAA and 30 µM PPBo treatments were associated with a decrease in grain weight at this time point (Fig. 2B). Already at 10 DAP, we observed an increase in unfilled caryopses in panicles treated with higher concentrations of IAA, L-kynurenine and PPBo. We excluded these unfilled grains from determination of grain fresh weight and IAA content. As the grains were maturing, the presence of unfilled grains became more pronounced (Fig. 3 A-C). Upon closer inspection of these grains, they appeared to be arrested in growth, suggesting a post-fertilisation seed abortion (Fig. 3C). We therefore counted the percentage of filled and unfilled/aborted grains in mature panicles harvested at 35 DAP. Indeed, compared to the control, all treatments except 100 µM L-Kyn were associated with high percentages of aborted grains, with the highest percentage (∼43%) recorded in treatments with 200 µM IAA and 500 µM L-Kyn (Fig. 2C). Furthermore, we found a significant negative correlation between percentage of aborted grains and grain IAA content at 10 DAP (Fig. 2D). Similar trends were also found in dry weight of whole panicles (Fig. 2E) and dry weight of 10 superior grains (Fig. 2G). All treatments except 50 µM IAA showed significant reductions in total panicle weight (Fig. 2E); a strong positive correlation was found between panicle dry weight and grain IAA content at 10 DAP (Fig. 2F). Consistent with the percentage of aborted grains, treatments with 200 µM IAA and 500 µM L-Kyn were associated with the largest reduction in dry weight of whole panicles (Fig. 2E) and 10 superior grains (Fig. 2G). However, in spite of high percentage of aborted seeds and reduction in total panicle weight (Fig. 2C and E), both treatments of PPBo did not show any significant change in grain dry weight compared to control (Fig. 2G). Furthermore, the correlation was also not significant between grain dry weight and grain IAA content at 10 DAP (Fig. 2H).

**Fig. 2.**
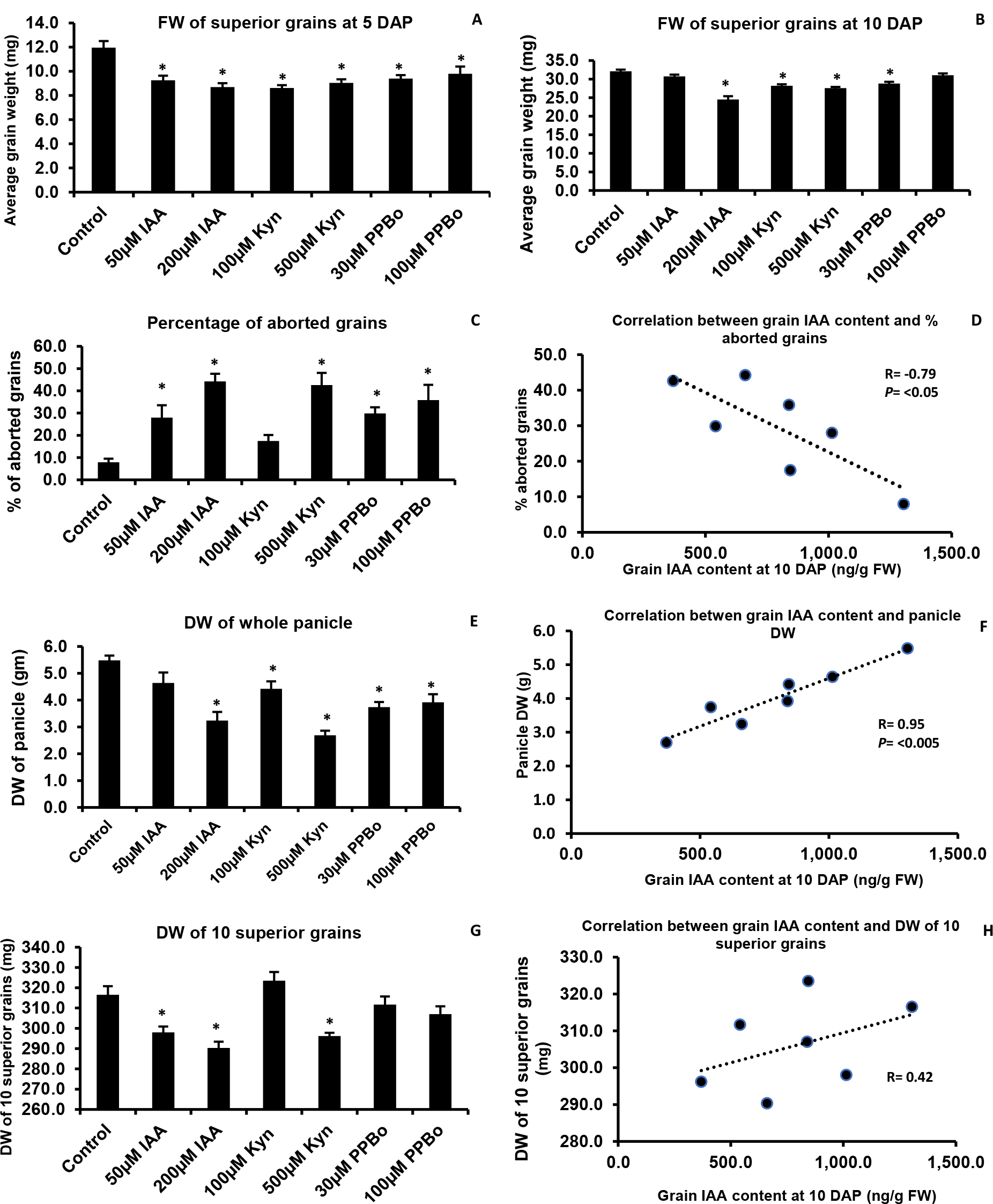
**(A-H)**. Yield-related parameters of chemical-treated and untreated immature and mature grains. Following application of IAA and IAA biosynthesis inhibitors (L-kynurenine and PPBo) to spikelets daily from 3 to 10 DAP, panicles were harvested at 5, 10 and 35 DAP (at maturity). Mature panicles were dried in an oven at 50^°^C for a week before measuring the yield parameters. Each data point in figures (A-H) represents the mean of six biological replicates ± the standard error of the mean (SEM). Data were subjected to one-way analysis of variance (ANOVA). In case of significant difference, mean values were compared using Tukey’s honestly significant difference (HSD) test. Asterisk denotes significant difference of a treatment when compared to the control at *P* <0.05. FW= fresh weight; DW= dry weight; DAP= days after pollination.

**Fig. 3.**
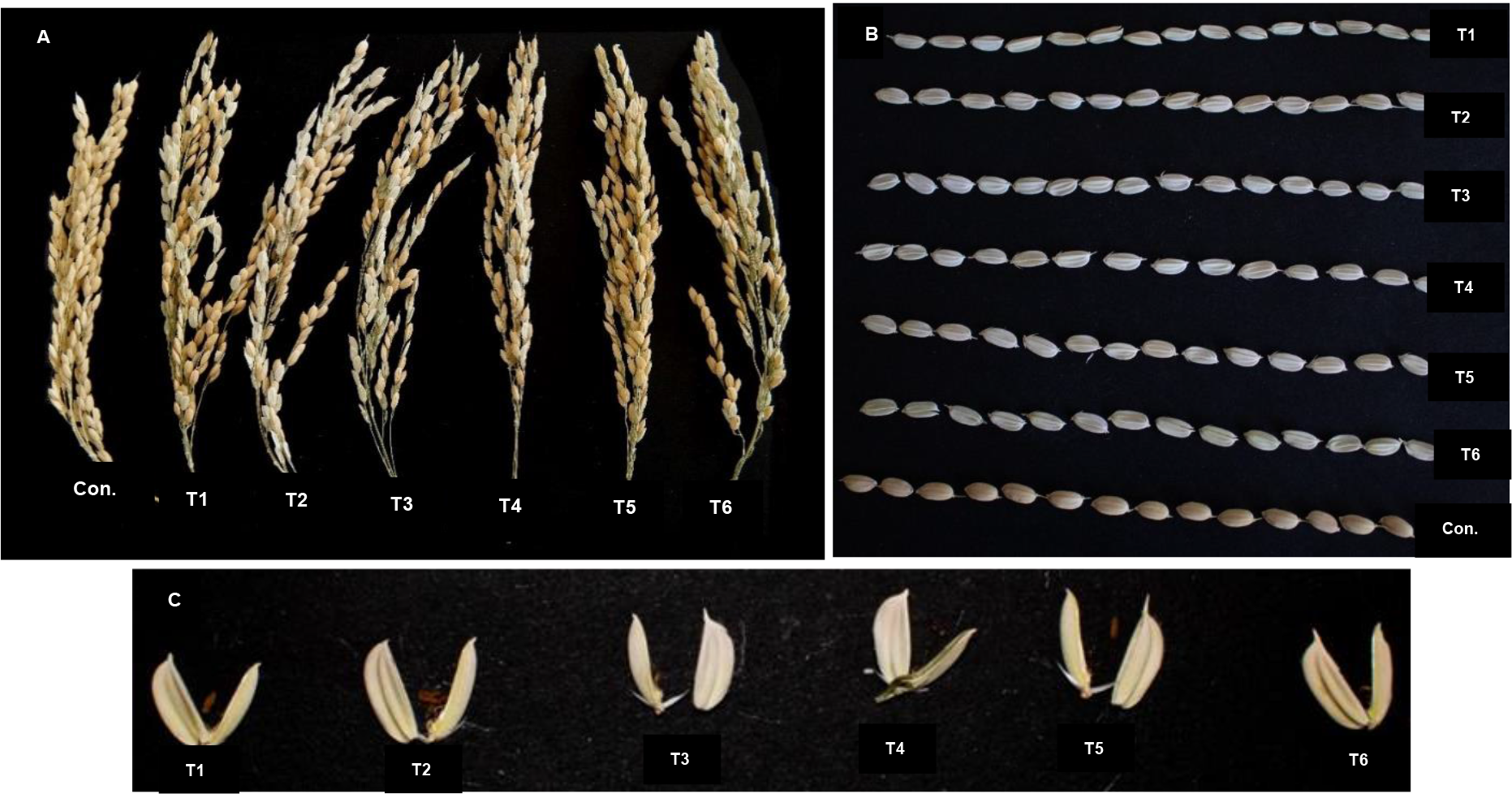
**(A-C)**. Images of mature panicles and grains that received chemical treatments during early stages of grain development. The panicles received chemical treatments from 3 to 10 DAP. They were harvested at 35 DAP (at maturity). (A) shows treated and untreated panicles at maturity; (B) shows unfilled/aborted superior grains from the treated panicles and compares them with the control; (C) shows individual unfilled/aborted superior grains from the six treatments at maturity. Con.= control (0.1% DMSO and 0.01% Tween-20), T1= 50 µM IAA, T2= 200 µM IAA, T3= 100 µM L-kynurenine, T4= 500 µM L-kynurenine, T5= 30 µM PPBo and T6= 100 µM PPBo.

## Discussion

Chemical biology is a powerful alternative to genetic and genomic techniques for investigating biological processes (Dejonghe and Russinova 2017). In this study, we used this approach to inhibit IAA biosynthesis in early rice grains and assessed the effects of this inhibition on final grain yield. We showed that both L-Kyn and PPBo were able to reduce significantly endogenous IAA levels in developing rice grains. The efficacy of these inhibitors has not been tested previously on cereal reproductive system. Our results thus show the potential of L-Kyn and PPBo in elucidating auxin biosynthesis and signal transduction during rice grain development. We also applied IAA to monitor its effect on grain development. However, grains treated with exogenous IAA showed a considerable reduction in endogenous IAA levels. IAA is known to stimulate its own metabolism. *GH3* genes encoding IAA-amido synthetases are concomitantly up-regulated in vegetative tissues with high IAA levels; excess IAA can also be sequestered as IAA-ester conjugates (Ludwig-Müller et al. 2009, 2011). Phenotypic effects of exogenous IAA application on yield parameters need therefore to be interpreted in light of actual endogenous IAA levels in IAA-treated grains.

Reduction in grain IAA levels appeared to be detrimental to grain filling. Our results are in agreement with several reports that provided correlational and genetic evidence for a crucial role of IAA in regulating starch accumulation in cereal endosperm (e.g. Abu-Zaitoon et al. 2012; Bernardi et al. 2012; Xu et al. 2021). Furthermore, Tamaki et al. 2015 reported a reduction in grain yield following application of the IAA biosynthesis inhibitor L-amino-oxyphenylpropionic acid (L-AOPP). However, unlike Ghorbani Javid et al. 2011 who recorded a positive effect of exogenous auxin application on grain-fill in rice, similar dosage of IAA used in this study failed to show any significant increase in grain weight. We interpret this in light of the reduced endogenous IAA pool of these grains during their early development stages.

Extensive post-fertilisation seed abortion following application of IAA and its biosynthesis inhibitors was an interesting observation in our study. This pointed to the importance of a critical threshold level of grain IAA during its early stages of development. Indeed, Guo et al. 2020b showed that single mutants of the auxin signal perception components *OsTIR1* and *OsAFB2*, viz. *Ostir1* and *Osafb2*, were characterized by reduced number of filled grains per panicle; this phenotype became more severe in case of *Ostir1Osafb2* double mutants. We chose to start inhibitor application at 3 DAP, because *OsYUC12* transcripts were first detected at this time (Basunia et al. 2021). Interestingly, formation of endosperm coenocyte becomes complete and cellularisation of coenocyte nuclei initiates at 3 DAP; cellularisation is complete by 5 DAP, by which time *OsYUC12* has peak expression in the entire aleurone (Wu et al. 2016; Basunia et al. 2021). The timing and progression of cellularisation are critical for endosperm development, as precocious and delayed cellularisation both affect the final grain size and may even result in seed abortion in rice as well as in *Arabidopsis* (Stoute et al. 2012; Folsom et al. 2014). Furthermore, a critical level of endogenous IAA is required for endosperm cellularisation to proceed in *Arabidopsis* (Batista et al. 2019). Disturbing this level by either reducing or elevating endogenous IAA may have inhibitory effects on cellularisation that could lead to seed abortion. The same mechanism may also operate in rice (Paul et al. 2020). It is probable that the grains which were not aborted and hence grew to maturity had the threshold IAA level, which allowed timely progression and completion of coenocyte cellularisation, and ensured consequently their retention. Conversely, grain retention may have occurred when the inhibitors did not penetrate into some ovaries, which consequently were able to produce sufficient IAA for normal endosperm development to proceed. This view is supported by the observation that, in spite of extensive seed abortion and subsequent reduction in total panicle weight, dry weight of PPBo-treated grains was similar to that of control grains. Overall, our results suggest that, besides regulating the grain-fill, auxin action may also be involved in earlier cellular events during endosperm development in rice.

In conclusion, we have been able to inhibit IAA biosynthesis specifically in early rice grains by a chemical biology approach. The auxin-deficient grains suggested a crucial importance of IAA in grain retention. Compared to genetic techniques, this approach can be cost- and time-effective by inducing a phenotype in a rapid, reversible and conditional manner as well as circumventing the problem of lethality and gene redundancy.

## Author contribution statement

HMN and MAB conceived and designed the experiments. MAB conducted the experiments with assistance from HMN, DB and MM. MAB and HMN analysed the data. MAB wrote the initial manuscript which other authors read, amended and approved.

## Acknowledgements

Rice seeds were kindly provided by the Yanco Agricultural Institute. MAB is grateful to University of New England for a post-graduate scholarship. The authors thank Michael Faint for his assistance with the greenhouse. This work was supported by internal funding from University of New England.

